# Dependency Map correlation analysis reveals WDR89 as a genome maintenance factor

**DOI:** 10.64898/2026.01.28.698016

**Authors:** Sean W. Minaker, Gian L. Negri, Gregg B. Morin, Peter C. Stirling

## Abstract

The Cancer Dependency map project (DepMap) has created an unprecedented resource of siRNA and CRISPR genetic screens across more than 1000 cell line models. Numerous computational tools have been developed to analyze DepMap outputs and have revealed new genetic dependencies relevant to cancer. Here we extracted correlation information from the DepMap using curated gene sets of biological pathways to determine if this approach could reveal new high confidence candidates. Beginning with a published set of curated DNA repair proteins we determined correlations of query gene knockout fitness across all DepMap cell lines. As expected, this approach extracted many known genome stability genes and also suggested several additional candidates not previously linked to genome maintenance. To validate our analysis we used siRNA depletion for candidate genes and identified several factors whose depletion increases DNA damage or delay repair. Among these was WDR89. WDR89 relocalizes to the nucleolus in an ATM dependent manner upon DNA damage and physically interacts with a number of nucleolar proteins. Depletion of WDR89 also abrogates nuclear p53 accumulation upon irradiation, suggesting a role in p53 signaling. These data establish WDR89 as a new genome stability regulator and highlight the value of DepMap for predicting previously unknown biology.

## INTRODUCTION

Genetic interactions define biological relationships between genes and pathways in cells. Quantitative fitness landscapes and digenic gene interaction profiling has been conducted systematically in several organisms, but none so completely as *S. cerevisiae* where a complete map of genetic interactions between all essential and non-essential genes is available (1–3). The goal of such profiling is to understand the wiring diagram of cell biology. This information has many benefits. One such benefit is to infer potential genetic dependencies in human diseases characterized by mutations in the genome, and thus suggest new precision medicine targets through a concept called Synthetic Lethality (4). Ideal genetic networks are generated in otherwise isogenic cells with known gene mutations or knockouts. In addition, genetic networks are conditional, changing with the environment, and epigenetic state of the cell population (1, 5). Thus, only some synthetic lethal interactions will be robust across cell types and disease states, while others will be highly specific to the system measured.

The Cancer Dependency Map project (DepMap) combines deep cancer cell line profiling, with genetic dependency screens, and chemical screens to create a map designed to identify therapeutic targets and biomarkers(6). This effort has been incredibly successful already, and has much more to teach us as it continues to expand. Since each cancer cell line query contains a different constellation of mutations, a unique tissue of origin and associated gene expression state, and are growing under lab conditions, the data does not produce a gene-gene interaction network directly as produced in model systems. Instead, DepMap reveals a fingerprint of fitness effects across hundreds of diverse cell lines. Biological functions are embedded in this data and can be observed within the correlations of gene knockout effects, for example genes within protein complexes should tend to have similar patterns of cell line dependencies in the DepMap (6). In addition, genes of unknown function should exhibit cell line dependencies that cluster with genes of similar function. If the known gene functional data are well annotated, then it should be possible to predict cellular processes in which unknown genes function.

Here we extract cell line dependency profiles for a curated set of genome stability maintenance and DNA repair genes and identify all factors with similar profiles but lacking a clear genome stability annotation. Direct testing by siRNA knockdown confirmed six candidate genes whose depletion increases spontaneous DNA damage or delays repair of radiation induced damage. While several of the identified factors do have emerging or newly established links to genome stability, WDR89 is an evolutionarily conserved but nearly completely uncharacterized protein. Here we report the first study of WDR89 physical interactions, localization, and phenotypic analysis of WDR89 depletion. Our data finds that WDR89 depletion leads to spontaneous DNA damage and delayed repair of ionizing radiation (IR) damage. WDR89 accumulates in nucleoli and physically interacts by immunoprecipitation with chromatin modifiers, and nucleolar proteins that have connections to p53 regulation. WDR89 depletion impairs the IR-induced relocalization of p53 to the nucleus. Together our work describes for the first time WDR89 as a protein with a role in scaffolding nucleolar chromatin associated proteins playing roles in cellular responses to genotoxins in the p53 pathway.

## MATERIALS AND METHODS

### Querying the Cancer Dependency Map

Gene dependency data for 990 cell lines were obtained from CRISPR-Cas9 genome-scale knockout library, generated as part of Broad Institute’s Cancer Dependency Map project (DepMap 21Q2, https://depmap.org/portal/download/). Pairwise spearman’s correlation was calculated between 17645 gene pairs. A manually curated list of 186 genes with a DNA repair related function (**Table S1**) (7–9) was used to select the top 20 genes showing the highest correlation to each of the members of the DNA repair set. Selected genes showing high correlation (top 20) with at least 5 DNA repair genes were prioritized for follow up analyses.

### Cell culture and siRNA knockdown

U2OS (ATCC) and HCT116 cells were grown in McCoy’s 5A medium, HEK293T cells were grown in Dulbecco’s modified Eagle’s medium (DMEM), with all media supplemented with 10% fetal bovine serum. siRNA treatments were performed using Qiagen Flexitube siRNAs at 50 nM concentration with RNAiMax transfection reagent for 72 hours before fixation and/or protein harvest.

### Immunoprecipitation

HEK293T cells were transfected with plasmids containing a WDR89-V5 construct under control of a CMV promoter using polyethylenimine-mediated transient transfection. 10 ug of plasmid was transfected to each 10 cm dish followed by a 48h incubation period to allow for protein expression. Cells were exposed to 5 Gy of x-rays and allowed to recover for 1 hour before protein extraction in NP-40 lysis buffer (50mM Tris-HCl pH 7.4, 150mM NaCl, 1%NP-40 and 5mM EDTA.)containing benzonase, protease inhibitors and phosphatase inhibitors. Lysates were cleared by centrifugation and incubated with anti-V5 antibody (Bethyl), anti-WDR89 (Bethyl) or control IgG bound to Bio-Rad Surebeads at room temperature for 1 hour. The beads were then washed with lysis buffer 3x followed by elution in 1x SDS-PAGE buffer (Bio-Rad). Samples were then subjected to mass spectrometry analysis or western blotting.

### Mass spectrometry

0.5 μL of 1M dithiothreitol (DTT) in HPLC water was added to the eluted protein and samples were incubated for 30 min at 60 °C with shaking at 1000 rpm. Samples were cooled to room temperature and 1.5 μL of 1M iodoacetamide (IAA) in HPLC water was added and the reaction was incubated for 30 min at room temperature in the dark. An additional 1 μL of DTT was added to quench the reaction. Single Pot Solid-Phase Enhanced Sample Preparation (SP3) was performed as previously published (10). Following SP3 treatment, analysis of peptides was carried out on an Orbitrap Fusion Tribrid MS platform (ThermoFisher Scientific) by data dependent acquisition as previously described (11).

### Mass spectrometry data analysis

MS raw data were searched in Proteome Discoverer suite (v.2.4.0.305, Thermo Fisher Scientific) against UniProt human reference database (20351 sequences, 2021/07/16). Precursor and fragment ion tolerance were set to 20 ppm and 0.6 Da, respectively. Dynamic modifications included Oxidation (+15.995 Da, M), Acetylation (+42.011 Da, N-Term), and static modification included Carbamidomethyl (+57.021 Da, C). Enzymatic cleavage rule was set at full trypsin, with a minimum peptide length of six. Up to two missed cleavages were allowed. Peptide-spectrum matches (PSMs) were validated with Percolator, where only PSMs with false discovery rate (FDR) < 0.01 were retained in the analysis. The precursor abundances from PSMs that map to unique proteins were aggregated to represent the protein abundance. MS data are available via ProteomeXchange with identifier PXD073023.

### Western blotting

Lysates were boiled in SDS-PAGE loading buffer and loaded onto 10% acrylamide SDS-PAGE gels for electrophoresis and subsequent transfer to 0.45 micron nitrocellulose membrane. Membranes were blocked in either 5% milk or 3% BSA in TBS-T buffer. Membranes were incubated overnight with primary antibodies at 4℃. Membranes were washed 3x in TBST followed by incubation with HRP-conjugated secondary antibodies at room temperature for 1 hour. Following a second round of washing, membranes were treated with ECL reagent (12) and imaging using a Bio-Rad Chemidoc imaging system.

### Immunofluorescence and microscopy analysis

U2OS cells grown on coverslips were transfected with siRNAs (or not) for 48 hours followed by irradiation with 2.5 or 5 Gy of X-rays depending on the experiment. Control cells were fixed 2 hours post irradiation while the remaining cells were allowed to recover for 24 hours prior to paraformaldehyde fixation. For 5 Gy experiments treatment was followed by recovery at 37℃ for the indicated times. Drug treatments included 10 µg/mL bleomycin for 1 hr, 10 µM camptothecin for 4 hr, 1 µM pladienolide B for 4 hr and 2 mM hydroxyurea for 2 hours. For ATM inhibitor experiments, cells were pre-treated with 10 µM KU-55933 for 1 hour followed by irradiation with 5 Gy of X-rays and then fixation and immunofluorescence staining as for siRNA experiments below.

Immunofluorescence staining was performed using anti-γ-H2AX antibodies (Abcam, ab81299, Santa Cruz, sc-517348), an anti-Nucleolin antibody (Novus, NBP2-44610), anti WDR89 antibody (Bethyl, A301-872A) or p53 antibody (Cell Signaling, #9282) at 1:250 dilution and Alexafluor 568 goat anti-rabbit secondary and Alexafluor 488 goat anti-mouse antibodies (Invitrogen) at 1:400 dilution. Samples were imaged at 100x using a Leica Dmi8 fluorescence microscope and Metamorph Imaging software. γ-H2AX foci were scored and analyzed using ImageJ and Graphpad Prism software, respectively. For experiments measuring nuclear intensity of p53 or WDR89, a nuclear mask based on DAPI staining was used to define regions of interest for intensity measurement in ImageJ (13).

### Proximity Ligation Assay

U2OS cells grown on 12 mm coverslips at 50% confluency were transfected with plasmids containing WDR89-V5 under control of a CMV promoter using Lipofectamine 3000 reagent “(ThermoFisher Scientific). 48 hours post transfection, cells were fixed in 4% paraformaldehyde for 20 minutes and permeabilized with 0.1% Triton-X 100 in PBS for 30 minutes. The permeabilized cells were incubated with overnight at 4°C with mouse anti-V5 antibody (Thermo Fisher, MA5-15253) and rabbit antibodies against KAP1 (Proteintech,15202-1-AP), WHSC1 (Proteintech, 22722-1-AP), CBX2 (Proteintech,15579-1-AP), CBX3 (Proteintech, 66446-1-Ig), NOC2L (Proteintech, 28509-1-AP), NPM (Proteintech, 10306-1-AP) and PHF2 (Bethyl, A303-457A-T). All antibodies were diluted 1:1000 in Duolink antibody buffer. Following overnight incubation with antibodies, all subsequence steps in the proximity ligation assay were according to the manufacturer’s protocol. Imaging and analysis was as previously described for immunofluorescence samples.

### Chromatin Fractionation

HCT116 cells were grown on 10 cm dishes to approximately 80% confluency and then subcellular fractionation was conducted according to a protocol from Abcam. Briefly, cells were collected and incubated for 15 minutes on ice in 500 µL of fractionation buffer (20 mM HEPES pH 74, 10 mM KCl, 2 mM MgCl_2_, 1 mM EDTA, 1 mM EGTA). The cells were lysed by passing through a 27-gauge needle 10 times followed by 20 minutes on ice. A sample of whole cell extract was retained prior to a subsequent spin at 720x g for 5 minutes. The supernatant containing the cytoplasmic fraction was removed and retained and the nuclear pellet was resuspended in 500 µL of fractionation buffer and passed through a 25-gauge needle 10 x followed by another spin at 720x g for five minutes. The pellet containing the chromatin was resuspended in TBS + 0.1% SDS and the chromatin was digested by adding 250 units of benzonase. Whole cell extract, chromatin and cytoplasmic fractions were used for western blotting as described above. Blots were probed with rabbit anti-WDR89 antibody, anti-GAPDH antibody (Invitrogen, #MA5-15738) and anti-H2AX antibody(Santa Cruz, sc-517336).

### Cell viability assays

U2OS cells were treated with 100 nM siRNAs against WDR89 or a scramble control. 24 hours post transfection, cells were harvested and seeded into 96 well plates which were irradiated and then allowed to grow for 72 hours. Viable cell counts were obtained using a CellTitre-Glo 2.0 kit (Promega) according to the manufacturer instructions and luminescence was measured using a Tecan Spark plate reader.

## RESULTS AND DISCUSSION

### DepMap correlation analysis identifies known and novel genome maintenance factors

Genes with similar genetic interaction profiles cluster together in pairwise genetic interaction screens from yeast, and have been used to infer novel gene functions (2). The DepMap provides a similar phenotypic fingerprint of gene perturbation effects across many cancer cell lines, highlighting most similar profiles on its web interface. We reasoned that curating a set of biological related gene queries and extracting genes that correlate repeatedly with curated queries gives us a chance to identify new biological functions for those genes. We extracted the top 20 correlated genes for a curated set of 186 genes that encode different types of DNA repair proteins. Our analysis generated a list of genes ranked by the number of query DNA repair genes that had a significantly similar DepMap dependency pattern (**Figure 1A** and **Table S2**). 127 genes were found within the top 20 correlations of a DNA repair gene from our query set. As our query list was not exhaustive, many of the top 127 genes we identified had well-defined roles in DNA repair or genome replication and maintenance. Additionally, regulators of cell death and cell cycle progression were represented among the top hits (**Figure 1B**). We reasoned that some poorly annotated genes within the list should have functions in DNA repair based on their correlations. Of the remaining genes, we chose seven candidates with a variety of molecular functions and varying degrees of literature linking them to DNA replication and repair.

**Figure 1.**
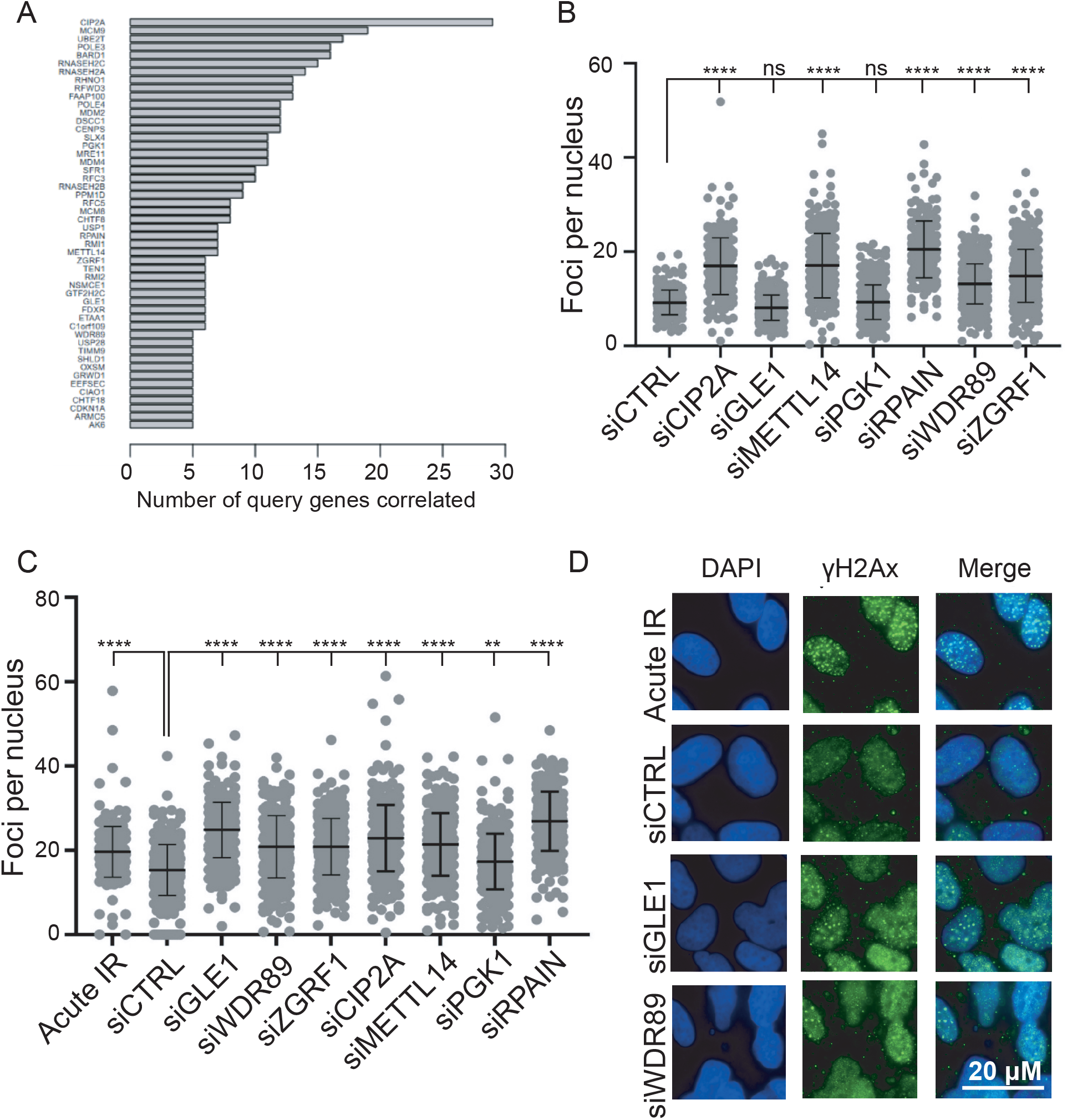
Extracting candidate genome maintenance genes from Cancer Dependency Map data. (A) Results for genes recurrently (>5 times) in the top 20 correlated dependency profiles for a curated set of DNA repair and genome maintenance factors. (B) γH2AX foci counts for U2OS cells treated with the indicated siRNA and imaged after 48 hours. (C) γH2AX foci counts for U2OS cells treated with the indicated siRNA, irradiated, and then allowed to recover for 24 hours. The first column, IR control, indicates foci numbers 2 hours after irradiation. ****p<0.0001, ANOVA with Tukey’s post-hoc test. (D) Representative images of γH2AX foci following irradiation and recovery in C. The top row shows no recovery period, compared to a 24 hour recovery for the indicated siRNA treatments below.

Under ambient conditions, knockdown of these seven genes resulted in significantly elevated ɣH2AX foci for the majority of genes tested (**Figure 1C)** indicating elevated levels of DNA damage. In order to assess whether depletion of these genes affected DNA repair, cells were exposed to 2.5 Gy of x-rays after 48 hours of knockdown and then stained for γH2AX foci 24 hours post-irradiation (**Figure 1D**). Strikingly, all the siRNAs tested resulted in a significant increase in γH2AX foci compared to the untreated control under some conditions (p<0.0001) (**Figure 1C**). Notably, siRNA treatment genes such as GLE1 and PGK1 did not show elevated DNA damage under ambient conditions but retained γH2AX foci 24 hour after irradiation, possibly due to direct or indirect defects in DNA repair. Depletion of CIP2A, METTL14 and RPAIN resulted in DNA damage foci comparable to the irradiated control two hours after x-ray exposure. Depletion of ZGRF1 and WDR89 led to more moderate induction of γH2AX both spontaneously and 24hrs after irradiation (**Figure 1C** and **1D**).

CIP2A, an important cancer biomarker (14), had a dependency map that correlated with the most query genes. CIP2A has now been identified as an important factor in genome stability through an interaction with TOPBP1 (15, 16), and regulation of nucleases and mitotic DNA synthesis (17, 18). Similarly, METTL14 has accumulated several links to genome stability in recent years (19, 20). RPAIN is poorly characterized but interacts with the well known single-stranded DNA binding protein RPA (21). ZGRF1 was described as a direct repair regulator of DNA end resection and D-loop stability (22, 23). Interestingly, WDR89 is completely uncharacterized as a genome maintenance factor. From this we can conclude that DepMap dependency profile correlations are a robust way to link a gene to a particular biological pathway, at least for DNA repair.

### Curated pathway level gene correlation extraction is useful in other biological processes

To determine if the approach of extracting multiple correlation ranks from a curated biological pathway list can generalizably identify components of that pathway we conducted three additional *in silico* experiments. We extracted correlated profiles based around gene lists for proteins associated with the 26S proteasome, the kinetochore and the nuclear pore complex. Again this approach successfully identified genes that were not included in the query list but that have well defined roles in these biological pathways (**Table S3**). The nuclear pore complex query revealed other NPC members, karyopherins and mRNA transport factors. The proteasome analysis revealed several additional proteasome subunits as well as numerous proteins without documented links to proteasome function while the kinetochore query set identified multiple proteins involved in kinetochore and microtubule functions. These data highlight the utility of Depmap analysis for identifying genes with connections to known biological pathways and protein complexes. For example, CLCC1 is a chloride channel that exhibits a dependency cluster with nuclear pore complex genes (**Table S3**). This result was initially puzzling but now a preprint has shown CLCC1 is the orthologue of yeast Brr6/Brl1 and regulates nuclear pore assembly (24). Similarly, c1orf109 has recently been implicated in chromosome stability through its interaction with the SPATA5-SPATA5L1 ATPase and we find it clustering with kinetochore components possibly due to effects on genome integrity (25). Thus, as with genome maintenance, other dependency clusters can be assembled and mined for novel associations to give first indications of function connections. Of course, testable predictions also emerge: CDCA3 is an E3 ubiquitin ligase linked to cell cycle regulation at mitotic progression in Xenopus and implicated in cancer (26–28). Based on this analysis correlating its dependencies with the nuclear pore we predict that CDCA3 may also have a function in regulating nuclear pores.

### WDR89 is a nuclear protein and accumulates in the nucleolus in response to DNA damage

While several of the top correlates from our genome stability analysis have been linked to DNA repair in the literature, the WD-40 repeat protein WDR89 is essentially uncharacterized in any context. Therefore, we chose to focus additional characterization on WDR89 to validate our approach for an unknown gene. In order to determine how WDR89 responds to DNA damage, we first performed immunofluorescence using an antibody against WDR89. WDR89 staining was seen in the nucleus and colocalized with nucleolin at the nucleolus (**Figure 2A)**. Nuclear/nucleolar intensity of WDR89 increased rapidly following an acute 5Gy irradiation, peaking within 60 minutes post-irradiation (**Figure 2B**). WDR89 levels increase in the nucleus both in response to irradiation and also when cells are treated with the radiomimetic chemical bleomycin (**Figure 2C**). However, other DNA damaging agents tested, hydroxyurea, camptothecin, or splicing inhibitor Pladienolide B, do not alter WDR89 staining intensity (**Figure 2D**).

**Figure 2.**
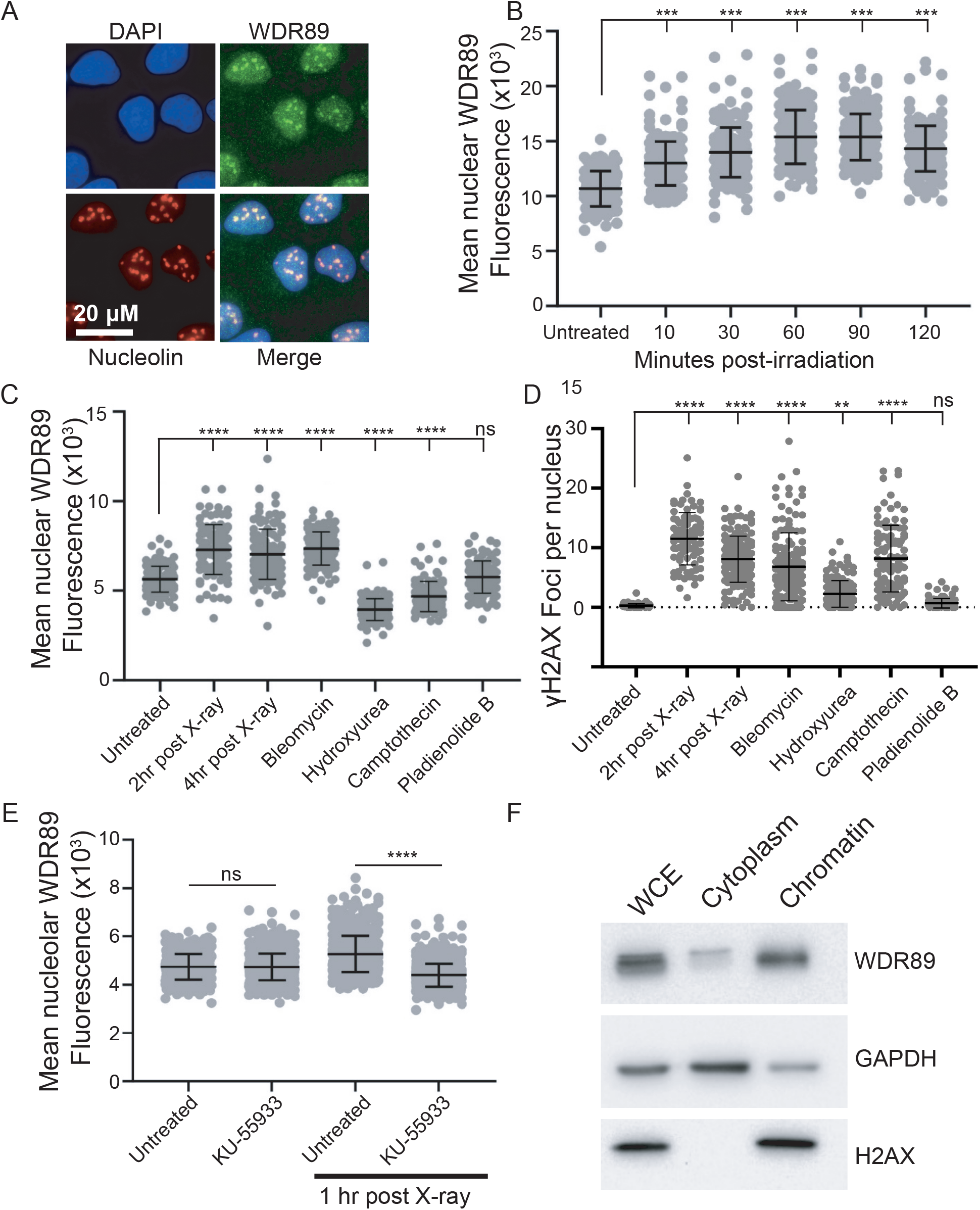
WDR89 is a nucleolar protein that responds to double-strand DNA break-inducing stresses. ATM-dependent nucleolar accumulation of WDR89 upon genotoxic stress. (A) WDR89 localizes predominantly in the nucleus and overlaps largely with nucleolin. (B) Quantification of WDR89 nuclear staining intensity following ionizing radiation. (C) Quantification of WDR89 nuclear staining intensity following genotoxic chemical treatments. (D) Confirmation of DNA damage by γH2AX foci staining of the conditions in C. (E) Effect of ATM inhibitor treatment on WDR89 nuclear intensity changes after IR. (F) Western blot of chromatin fractionation for WDR89 and the indicated markers. For panels in B, C, D and E, ****p<0.0001, by ANOVA with Tukey’s post-hoc test or by T-test where appropriate.

WDR89 has reported phosphorylation sites at S67, Y69, and S156 in high-throughput studies (29) and ATM signaling has been reported to regulate nucleolar chromatin organization in response to DNA damage causing double-strand breaks (30). We therefore decided to test whether treatment with agents that affect DSB signaling through ATM alter the dynamic localization of WDR89. We observed that pretreatment with the ATM inhibitor KU-55933 had no effect on baseline WDR89 localization, but prevented the increase in nucleolar WDR89 signal 60 minutes post-irradiation (**Figure 2E**). These results highlight a potential requirement for active DNA damage response signaling by ATM to regulate the accumulation of WDR89 in nucleoli during the DNA damage response. Finally, we suspected that WDR89 may reside in protein complexes associated with chromatin, so we conducted chromatin fractionation experiments on HCT116 cells and confirmed strong association of WDR89 signal by western blot with the chromatin fraction (**Figure 2F**).

### WDR89 protein interaction identify a network of chromatin and p53 regulators

To date, no studies have sought to define WDR89 protein-protein interactions so we used immunoprecipitation followed by mass spectrometry (IP-MS) to gain an initial picture of the protein-protein interactions for WDR89. Given the radio-sensitive localization of WDR89 we conducted V5-immunoprecipitation experiments in cells expressing WDR89-V5 with and without 5 Gy of radiation. Many of the top predicted interactors from our IP-MS included proteins associated with heterochromatin dynamics, chromatin modification and the nucleolus (**Table S4**). Many hits from mass spectrometry could be false positives, and indeed some such as NPM1 are frequent flyers in the CRAPome database (31).

In order to confirm at least some of the WDR89 IP-MS experiments, we used immunoprecipitation with two different antibodies, targeting either the V5-epitope tagged WDR89 or a native WDR89 antibody in irradiated cells (**Figure 3A**). Precipitation of WDR89 enriched WHSC1/NSD2, CBX2, PHF2, NPM1, CBX3, KAP1 and NOC2L. These proteins were chosen for validation because there was some evidence of multiple protein complex subunits in the WDR89 complex enrichment. All of these interactions could be detected with both native WDR89 antibody or with the anti-V5 tag antibody, except for CBX3 which only precipitated with the native WDR89 antibody (**Figure 3A**). It is possible that the V5 tag interferes with the CBX3 interaction directly or indirectly.

**Figure 3.**
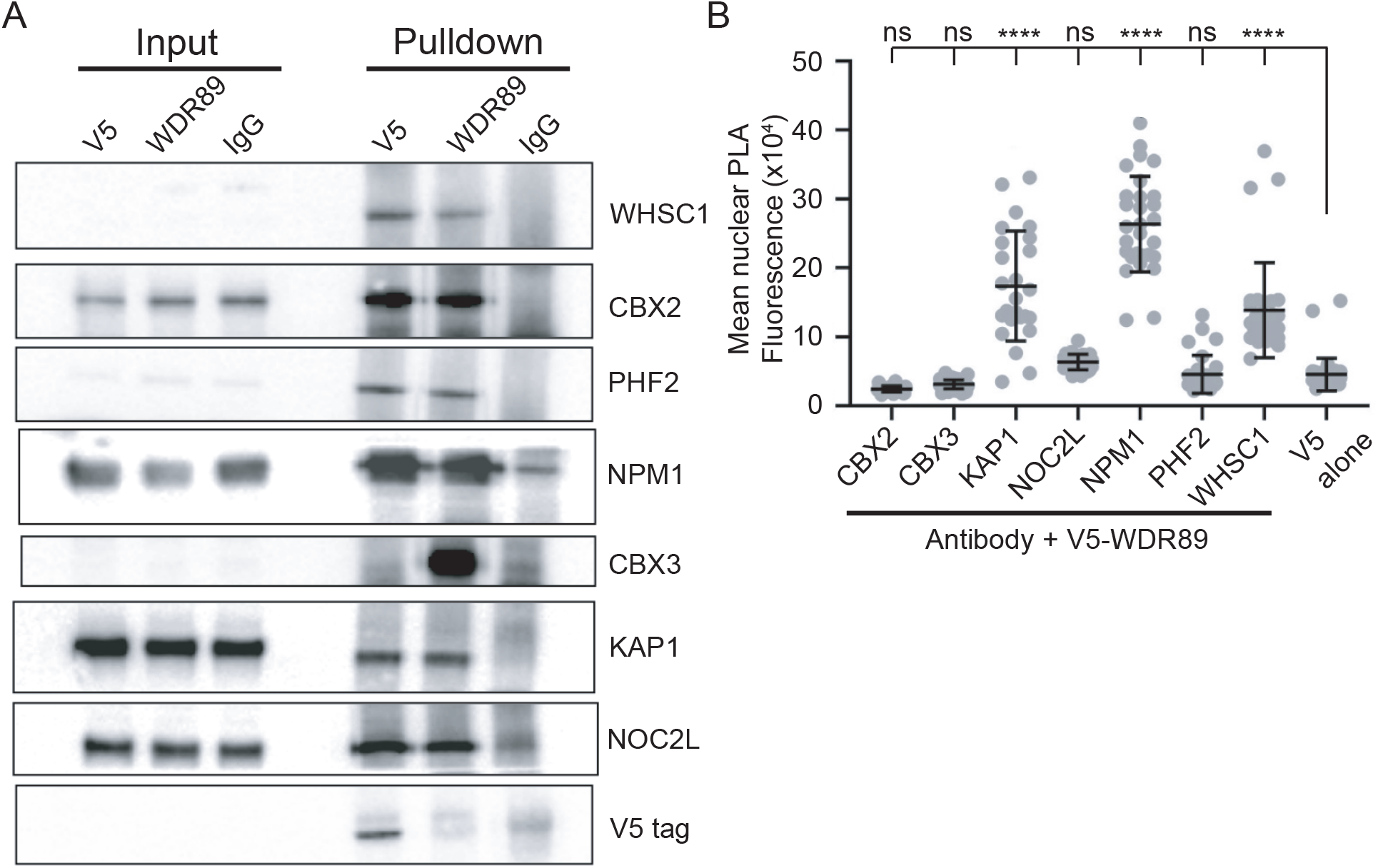
The WDR89 interactome includes nucleolar proteins and chromatin modifiers. (A) Western blot analysis of candidate WDR89 interacting proteins in V5-or native WDR89 pulldowns. (B) Proximity ligation assay for WDR89 with interacting candidate proteins. ****P<0.0001 by ANOVA with Tukey’s post-hoc test.

As a third orthogonal validation, we applied the proximity ligation assay (PLA) to observe WDR89 proximity with some of its potential interacting partners. KAP1/TRIM28, NPM1 and WHSC1 generated significant PLA signals with WDR89 compared to the V5 control, while the other partners did not (**Figure 3B**). Failure to generate PLA signal does not confirm lack of interaction as it can occur for a variety of reasons including the compatibility of the antibodies with the assay, however the positive signals for KAP1, NPM1, and WHSC1 strongly support the mass spectrometry and western blot data. Importantly, all of the WDR89 interactors are nuclear proteins, many of which have functions within heterochromatin or the nucleolus consistent with the observed localization of WDR89.

Functionally, the WDR89 interacting partners define a tightly focussed and coherent group of chromatin regulators placing WDR89 within this network for the first time. WHSC1, also called NSD2, is a histone methyltransferase that creates active H3K36me2 and a potential cancer drug target (32). PHF2 is a histone demethylase that functions with cohesin in gene regulation (33, 34). KAP1 binds to hypomodified H4 tails to promote RNAPII pause release (35). Chromobox protein CBX2 is a component of the H2AK119 ubiquitylating Polycomb Repressive Complex I, while CBX3 helps to retain heterochromatin at the nuclear lamina (36). NPM1 is a cancer gene whose encoded protein is dynamically nucleolar and impacts nucleolar structure (37). Finally, NOC2L, also called NIR, is a nucleolar and nucleoplasmic protein that inhibits histone acetyltransferases and exhibits physical interactions with both p53 and its regulator MDM2 (38, 39). Overall, these interactions place WDR89 as a partner for several chromatin modifying enzymes, with strong links to both nucleoli, cancer, and the p53 pathway during DNA damage.

### WDR89 depletion affects the p53 response

Revisiting the physical interactions of WDR89 in the context of the DepMap correlation data illustrates a compelling potential connection to the tumor suppressor p53. NPM1, KAP1, CBX2, and NOC2L all interact with p53 in the STRING database. Moreover, negative regulators of p53 such as MDM4, MDM2, and PPM1D are among the top positive co-dependencies of WDR89, while the top negative co-dependency of WDR89 is p53 itself (https://depmap.org/portal/gene/WDR89?tab=overview). A physical interaction between WDR89, NOC2L and the other partner proteins provides a potential explanation, suggesting that WDR89 may function to negatively regulate p53 responses. This in turn could potentially contribute to the DNA damage repair phenotypes observed.

To begin to test this association we first tested whether knockdown of WDR89 by siRNA affected p53 expression or localization in response to irradiation. **Figure 4A** shows a slight increase in p53 upon WDR89 knockdown in untreated cells and that ionizing radiation strongly induces p53 nuclear staining in control cells as expected. Remarkably, nuclear p53 localization does not increase in cells siRNA depleted for WDR89 and exposed to ionizing radiation (**Figure 4A** and **4B**). Conversely, WDR89 localization after irradiation was measured in HCT116 cells with or without p53 knockout. **Figure 4C** shows that, in p53-wildtype cells, WDR89 increases nuclear intensity after irradiation, while in p53-null cells WDR89 nuclear intensity actually drops slightly after irradiation. Failure to respond correctly to double strand breaks might ultimately lead to chromosome instability. Consistent with this hypothesis, siRNA depletion of WDR89 significantly increased the frequency of micronuclei in both U2OS and HCT116 cells compared to control siRNA (**Figure 4D**). Finally, if p53 responses are indeed muted in WDR89 depleted cells, then cell survival after irradiation could be impacted. We siRNA depleted WDR89 in U2OS and then did a dose-response of survival in irradiation (**Figure 4E**). This analysis showed that WDR89 depletion led to significantly more surviving cells, consistent with a weakened p53 response.

**Figure 4.**
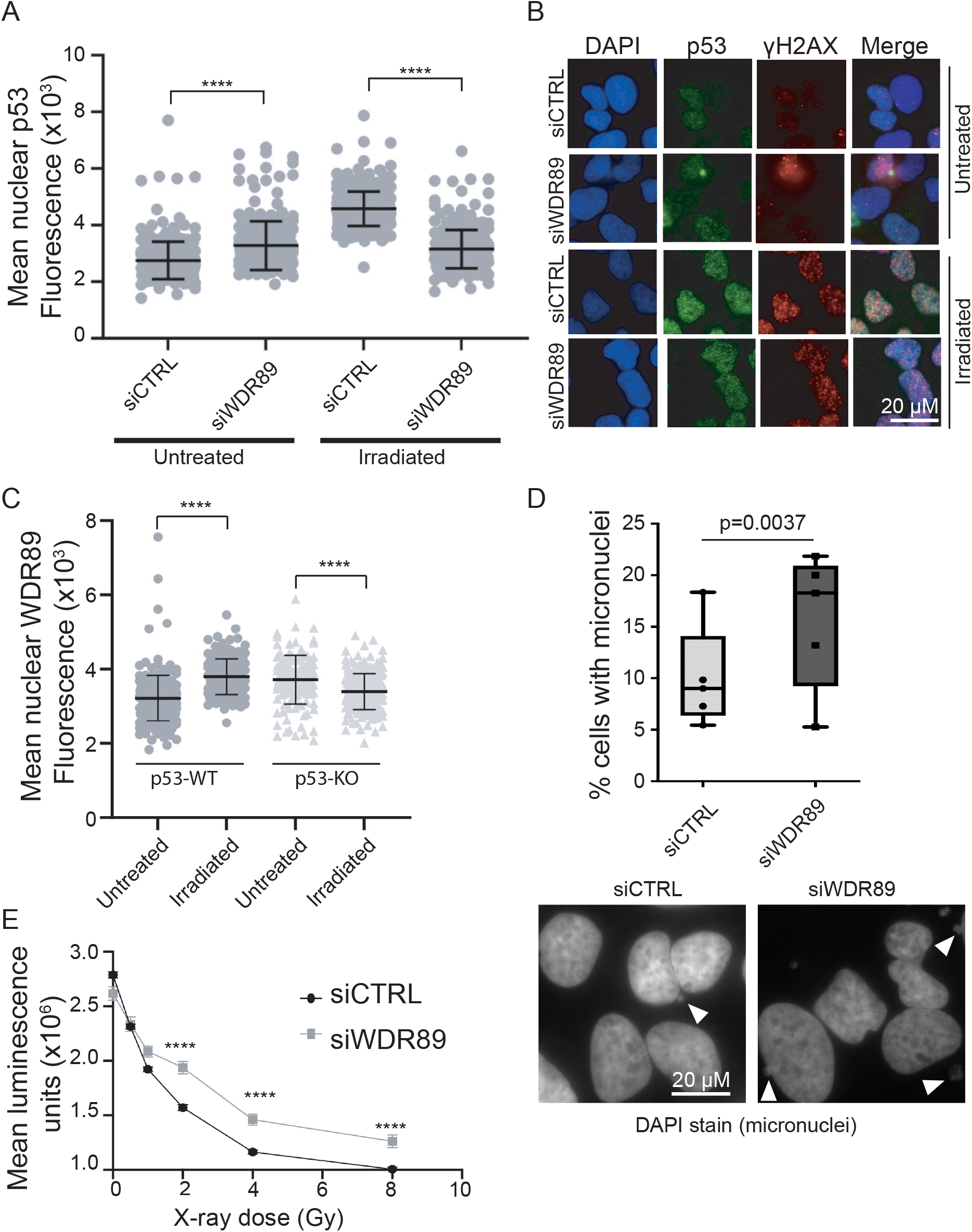
Inter-relation of p53 and WDR89 dynamics. (A-B) Irradiation induced nuclear p53 accumulation with the indicated siRNA treatment. A, quantification. B, representative image. (C) WDR89 nuclear accumulation in p53-WT or p53-KO cell lines. (D) Quantification of cells with an associated micronucleus in the indicated siRNA treatment. P-value determined by Fisher’s exact test of five pooled replicates. Top = quantification; bottom = representative image. (E) Ionizing radiation cell survival measurements with the indicated siRNA treatment. For A, C, E, ****p<0.0001 by T-test.

### Perspective

The Cancer Dependency Map CRISPR screening dataset is an unprecedented resource that has largely focussed on identifying synthetic lethal genetic liabilities that can be deployed in specific cancer types or against specific biomarkers (6). This is an outstanding goal whose success is being exemplified by the WRN-helicase inhibitors now in clinical trials for microsatellite unstable cancers (40). However, by creating a high resolution phenotypic fingerprint across diverse cancer cell lines, the DepMap data also has a massive potential to reveal novel biological functions for any human protein that has fitness impacts in cultured cells. Here we mine the predictive power of DepMap data to identify WDR89 as a genome maintenance factor with correlated dependency profiles to multiple genome stability factors. The top correlated single DepMap profile for WDR89 is DDX31. DDX31 itself is quite poorly characterized but was implicated as a partner of MDM2 in a 2012 paper, where its depletion promoted NPM1 binding to MDM2 (41). Indeed, DDX31 also interacts with NPM, further strengthening the ties to WDR89 (41). This aligns well with our analysis which shows that WDR89 interacts with p53 regulators like NPM1 and NOC2L and functionally impacts spontaneous DNA damage, DNA repair, and p53 localization in response to ionizing radiation. Our data suggest that WDR89 exerts its role in genome maintenance at least in part by interacting with chromatin and nucleolar organizing proteins that impact p53 signaling.

WDR89 is a conserved protein with a predicted beta propeller structure and no other functional domains. While it has unknown function across species, it is notable that the yeast homologue YNL035C (www.yeastgenome.org) has been annotated as increasing in abundance in response to DNA replication stress and has potentially emerging links to ribosome production and thus nucleoli (i.e. a reserved gene name annotates it as a chaperone for Ribosomal Protein 1; RPL1). Perhaps even though p53 is not conserved to yeast, there is some ancient shared function related to nucleolar DNA damage response signaling that involves the WDR89 family. Here we illustrate the value of DepMap correlations for functional annotation of unknown proteins like WDR89. Considerable additional work is needed to understand the mechanism and regulation of WDR89, as it pertains to p53, DNA repair, or other functions.

## ACKNOWLEDGMENTS

This work was funded by the Canadian Institutes of Health Research Project 398871 to P.C.S. We thank members of the Stirling lab for helpful discussions and the team at the BC Cancer Proteomics Platform for help with analysis.

## FIGURE LEGENDS

**Supplementary Table S1**. Query Genes used to extract common DepMap Correlates

**Supplementary Table S2**. Genes with recurrently correlated genome maintenance DepMap profiles.

**Supplementary Table S3**. Kinetochore, proteasome and nuclear pore DepMap extraction network.

**Supplementary Table S4**. Mass spectrometry results for WDR89 Immunoprecipitation experiments

## Notes

### Competing Interest Statement

The authors have declared no competing interest.

